# Mechanical Work Performance Constraints and Timing Govern Human Walking: A Modified Inverted Pendulum Model for Single Support

**DOI:** 10.64898/2026.03.09.710603

**Authors:** Seyed-Saleh Hosseini-Yazdi, John EA Bertram

**Affiliations:** Department of Biomedical Engineering, Schulich School of engineering, University of Calgary, AB, Canada; Human Performance Laboratory, Faculty of Kinesiology, University of Calgary, AB, Canada; Cumming School of Medicine, University of Calgary, AB, Canada

**Keywords:** Walking pendular model, Risk of fall, Step mechanical energy dissipations, Step work performance capacity, Hip joint mechanical work, Walking assistive device, Step-to-step transition, Single support mechanical work

## Abstract

Human walking is often considered an inverted pendulum during single support, suggesting conservative dynamics. Gait consists of discrete steps connected by mechanically costly transitions. We examine how step length, walking speed, and work capacity jointly constrain walking mechanics. Using a powered simple walking model, minimum speed required to complete a step of given length is derived based on gravitational work; below this threshold, forward progression becomes mechanically infeasible, and the next heel-strike occurs early, producing shorter steps. Comparisons with empirical step length–speed relationships show that humans walk at higher speeds and require greater push-off work, indicating energy dissipation. We extend pendular dynamics by incorporating hip torque, a linearized axial force model, and muscle intervention. This framework reproduces key GRF features, including the M-shaped profile, without prescribing force trajectories a priori. Fitted parameters suggest reduced average loading (*C*_*Baseline*_ < 1), active mid-stance unloading (*A*_*m*_ < 0), and narrowly timed muscle action (small *σ*_*m*_). Parameter studies show that increasing step length or speed increases transition work and peak forces, while hip torque timing indicates mechanical cost is minimized when energy modulation occurs after mid-stance. These findings indicate that preferred walking speed emerges from feasibility and work-capacity constraints, not energetic optimality alone.

## Introduction

Human walking is commonly described as resembling the motion of an inverted pendulum (1), especially during the single-support phase in which the center of mass (COM) vaults over the stance limb in a way that is largely considered conservative (2,3). With sufficient initial mechanical energy, such motion can, in principle, complete the step without additional work (4). However, this analogy is incomplete. Walking is not a single pendular motion, but a sequence of steps that requires repeated transitions from one supporting limb to the other (5). These step-to-step transitions incur substantial mechanical work and account for a large fraction of the energetic cost of walking (6). Subsequent to any step-transition, single support work may also regulate COM mechanical energy by performing net mechanical work (7–9). Although there is a simple mechanistic model to explain the step-to-step transition (5), there are limited comparable models for active work during single support.

The ability to perform adequate step-transition work is critically important to successful walking. When the preferred energetic pathway—pre-emptive push-off—is insufficient to offset the mechanical energy losses associated with collision-based losses, compensatory sources of positive work must be recruited (7). Consequently, active mechanical work is observed throughout stance, i.e. during both double (5) and single support (7), either to dissipate excess energy or to compensate for energetic deficits when walking conditions are imposed (8,9). Not surprisingly, since active work is continuously applied to regulate the COM state during stance (7), a purely passive pendular model cannot fully describe human walking and requires modification to account for these active contributions.

By definition, walking speed, step length, and step frequency are tightly interrelated (10). Empirical evidence suggests that walking speed and step length are largely determined by optimizing, as much as possible, to the cost surface in speed-frequency-cost space (11). This means that speed, step length and frequency will follow power-law relationships (10), such that constraining any two parameters determines the third through energetic constrained optimization (11). In experimental studies, walking conditions are typically prescribed, even if that is only in terms of speed (as in treadmill walking), prompting individuals to adjust the remaining parameters accordingly (6,12). In contrast, during unconstrained (free) walking, the factors that determine preferred gait parameters are less explicit and may reflect more fundamental mechanical constraints. With the advent of instrumented treadmills, experimental and simulation studies of walking have become predominantly speed-driven (5), implicitly treating walking speed as the primary control variable (10). Under this framework, step length, energetic cost, and mechanical work emerge as dependent outcomes. Here, we contend that this perspective overlooks a key constraint: the work required against gravity and the capacity to perform step transition work. Step length directly influences the required walking momentum and, in turn, the magnitude of step-to-step work necessary to maintain forward progression (5). Thus, a walker must be able to perform the required active work to provide walking speed sufficient to satisfy the mechanical work demands of successful single support completion imposed by the chosen step length (6,13).

Accordingly, we propose that for any prescribed step length, there exists a minimum walking speed required to complete the step based on pendular motion (required gravity work) (2). To maintain this required momentum, the inevitable energy losses at step transition must be compensated for (5,13). If the necessary work cannot be delivered through the pre-emptive push-off, mechanically less favorable strategies—namely post-transition single-support active work (rebound–preload)—must supply the remaining energy (Figure *1*) (9). Alternatively, the walker may adopt shorter steps, resulting in slower walking speeds (10,11). Therefore, there are a wide array of cases for which single support is not mechanically passive (8). Correspondingly, we will attempt to extend the classical pendular framework to incorporate both passive (elastic) and active single-support work.

**Figure 1.**
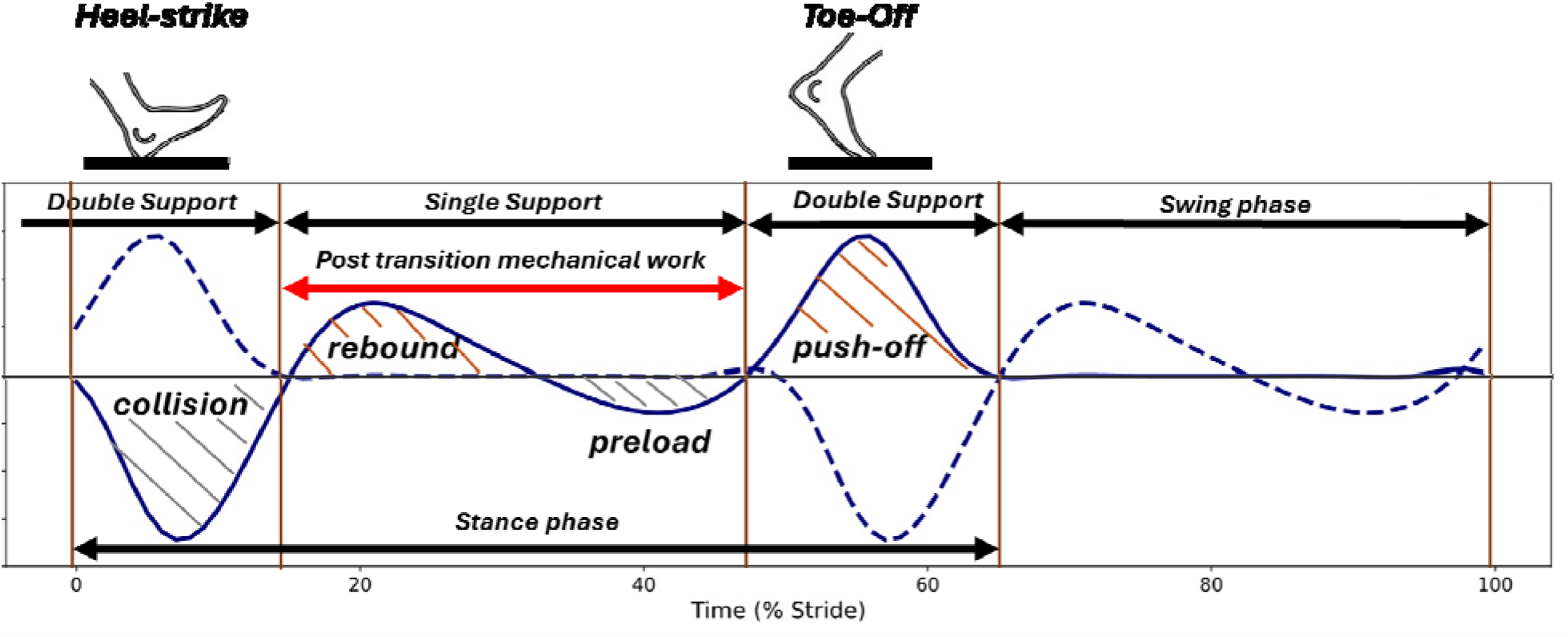
Based on GRF data, the Center of Mass (COM) power trajectory can be developed. It reveals two key instances in which humans perform positive work. The first instance occurs just before the step transition, when the trailing leg generates positive work, referred to as push-off. This work is primarily associated with the ankle. The second instance occurs during the single support phase and is known as rebound. Under normal walking conditions, the rebound is fully absorbed by the subsequent negative work (preload). However, if the rebound contributes to energizing the gait, its magnitude exceeds that of the preload, resulting in a net positive single-support mechanical work (adopted from *(7)*).

## Materials and Method

For our simulation, we use a powered simple walking model (5). This model assumes that the subject’s entire mass is concentrated at the Center of Mass (COM), with legs modeled as rigid and massless. We examine the motion of an inverted pendulum from three perspectives. (I) From a kinematic view, it is demonstrated that the equation of motion for an inverted pendulum can be represented as follows (5):

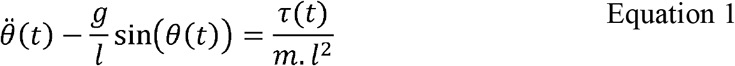

The angle ‘*θ*(*t*)’ is the instantaneous angle between the stance leg and vertical. Following the right-hand rule, the direction of positive angle and the angular velocity are shown in Figure 2A. Since the progression is from left to right, the direction of the angular velocity is negative. To solve the equation of motion, we assume the initial conditions as ‘2*α*’ that represents the leading and trailing leg angle at the double support phase (*α* =0.4), and 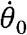for the initial velocity. Therefore, the initial linear velocity of the COM becomes: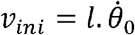, in which ‘*l*’ represents the leg length. Here, we consider the leg length as one. We solve the equation of motion numerically and end it when the instantaneous leg angle with the vertical becomes ‘− *α*’.

(II) From a kinetic viewpoint, we consider the initial position of the COM as the basis for the gravitational potential energy, i.e. datum or potential energy reference, thus; *E*_*p-initial*_ = 0. We assume the motion of the COM is of an inverted pendulum that is conservative and passive thus, *τ* (*t*) =0 (2,3,5). Having an initial velocity at the start, to reach mid-stance where the elevation of COM is maximum, the magnitude of the required work against gravity is:

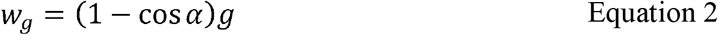

**Figure 2.**
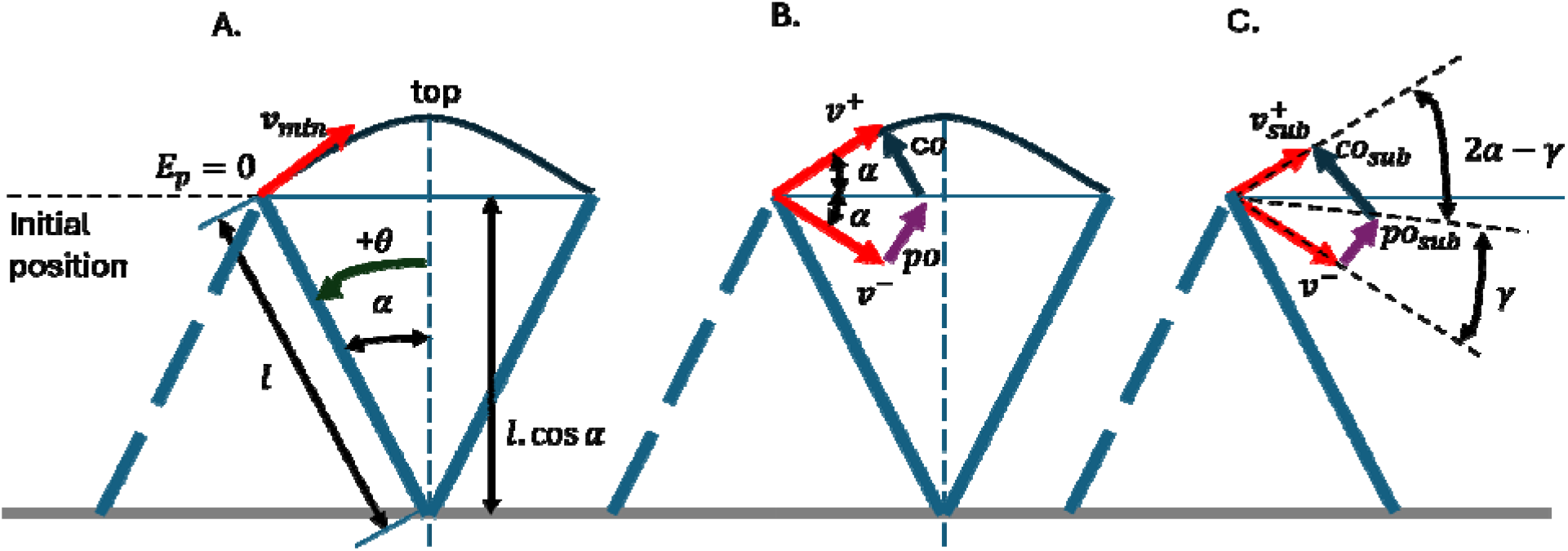
(A) The single support phase of human walking that resembles the motion of an inverted pendulum. Since the system is conservative, its total energy remains constant. Consequently, for a given step length, we can estimate the minimum require walking speed at the onset of single support. In this analysis, the counterclockwise direction is defined as positive direction for both angle and angular velocity. (B) To maintain forward progression, humans switch from one stance leg to the other. At the point of transition, there is a large mechanical energy dissipation that must be compensated for by active muscle work, ideally performed pre-emptively. (C) If the pre-emptive push-off is insufficient, the post-transition walking speed decreases. In such cases, additional post-transition energy compensation is necessary to sustain walking.

That represents the normalized work. Therefore, the minimum necessary initial COM velocity must be:

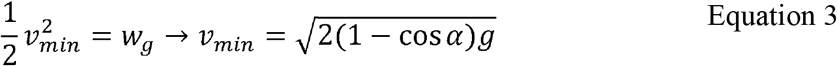

Therefore, for any given step length, we can find the minimum required initial speed of the pendulum phase. We call this the speed derived from the *gravity method*.

(III) On the other hand, human walking is not always pendulum-like, i.e. at the point of step transition (6,13) that coincides with mechanical energy loss (13). The preferred method of performing positive work required to compensate for this energy dissipation is the application of a pre-emptive push-off (5). Under nominal walking conditions, the magnitude of push-off matches the magnitude of the collision (energy loss) (13). Therefore, the push-off ensures the minimum COM velocity necessary at the start of the new stance (Figure 2B). For a given walking speed that satisfies the minimum required kinetic energy of prescribed step length (gravity method), the optimal push-off impulse is (5):

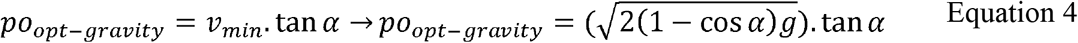

The push-off is calculated as 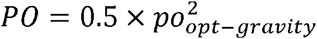(14). We consider the range of 0.64 m to 0.8 m for step length (2*α*) to evaluate the minimum required momentum and step positive work (push-off). Since there is an empirical power law relationship between step length and walking speed (10), we also derive the empirical value for walking speed for the prescribed step lengths. This allows the calculation of the analytical step-transition push-off which can be compared with the gravity work method result.

From the motion dynamics viewpoint, the axial force in the pendulum is 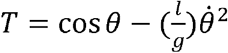(2,3). It is fair to assume that a human walker cannot maintain its angular velocity throughout the stance phase perfectly, and hence, we can expect the presence of some small perturbations. Thus,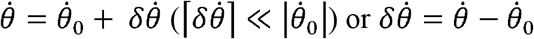Then:

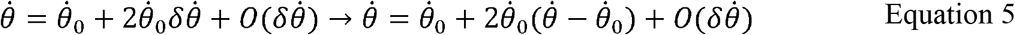

Therefore,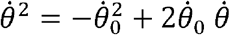. As such,

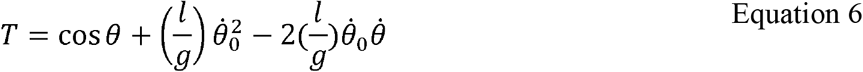

It means that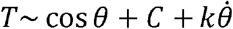, in which ‘*C*’ represents a baseline and ‘*k*’ is similar to a linear dissipation. Hence, we can replace the quadratic term with its first order approximation, and due to its representation, we call it a ***Linearly Damped Model*** just to distinguish it from the quadratic model. As a result, the vertical reaction force becomes *Fz* =*T*.cos*θ*. To account for the impact of muscle force application, we also add a muscle modulation factor as 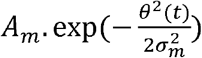This enforces the muscle term to be a mean of zero over the step so that force redistribution can be changed without changing the net impulse. Thus, the muscle intervention does not dissipate any energy. *A*_*m*_ is the amplitude of force redistribution (dimensionless body weight or BW). When *A*_*m*_ < 0 muscle unloads the limb at mid-stance, and when *A*_*m*_ > 0, muscle increases the mid-stance support, and a larger |*A*_*m*_| demonstrates a greater active control of vertical loading. The *γ*_*m*_ represents muscle modulation timing (width) to control angular speed of muscle action (radians). Thus, a small *σ*_*m*_ indicates a brief, sharply timed activation near mid-stance, while a large *σ*_*m*_ indicates a broader activation during the single support phase. Lastly, to adjust the vertical force trajectory baseline we adopt a proportionality coefficient *Fz*_-*Model*_ = *C*_*Baseline*_ *F*_*z*_(dimensionless). While *C*_*Baseline*_ < 1 shows partial loading, *C*_*Baseline*_ ∼1 indicates near full weight support.

The, *A*_*m*_ *σ*_*m*_ and *C*_*Baseline*_ are calculated as follows when we compare the average human walking vertical GRF, at S = 0.7m, and therefore, human walking speed of *v*_*ave*_ =1.2 m.s-1, with model force associated for the same step length (*α* =0.35 rad.):

We select a portion of vertical GRF that is suggested to be associated with the single support phase (15) for model training.

Then, we assess the resulting vertical reaction force peaks and trough for given step lengths (0.6 m, 0.7 m, and 0.8 m) and their associated speeds derived from the gravity method.

Thereafter, we run a parameter study to examine the impact of the adopted walking speed for a given step length on the vertical reaction force of the linearly damped model. We start the parameter study with, when . We then increase the walking speed.

We also quantify the impact of COM mechanical energy modulation through torque application (e.g. hip). This indicates how the vertical reaction force would change when COM energy i wasted or increased by application of opposing or supporting hip torques. Therefore, the initial velocity must be slightly larger than the minimum recommended by the gravity method to compensate for lost mechanical energy of opposing hip torque. For *α* = 0.4 *rad*, we consider initial velocity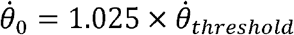. For simulation, the supporting and opposing torques are -0.15 (in the direction of motion), and 0.15 (opposing the direction of motion), respectively.

Human walking hip torque trajectories for different speeds indicate positive torque in early stance, while the torque becomes negative from midstance to late stance (16). Therefore, we consider the first vertical force peak, i.e. end of heel-strike double support (15), as the point where hip modulation strategy changes. We compare the push-off and collision peak differences when the hip torque is positive (opposing), zero, and negative (supporting) before the first vertical force peak. After this point, the hip torque is negative (supporting) until the end of double stance. We compare the hip net work for each case.

## Results

For *α* =0.4 rad., we considered three scenarios: less than, equal to, and greater than the threshold defined by Equation 3 (gravity method,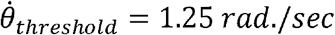). When the initial velocity is less than the threshold 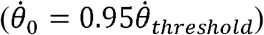, the COM progresses in the desired direction but eventually speed falls to zero, causing the COM to fall backward, as indicated by a positive angular velocity (Figure 3A). When the initial velocity is equal to the threshold, the COM reaches the mid-stance point with a speed of zero. This point represents an unstable equilibrium where a slight perturbation could push it in either direction (Figure 3B). Once the initial velocity exceeds the threshold magnitude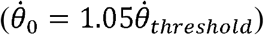, the COM successfully completes the step (Figure 3C). These findings indicate that the COM’s initial velocity substantially affects gait stability in the sagittal direction. Consequently, the inability to provide the required walking momentum results in a fall.

**Figure 3.**
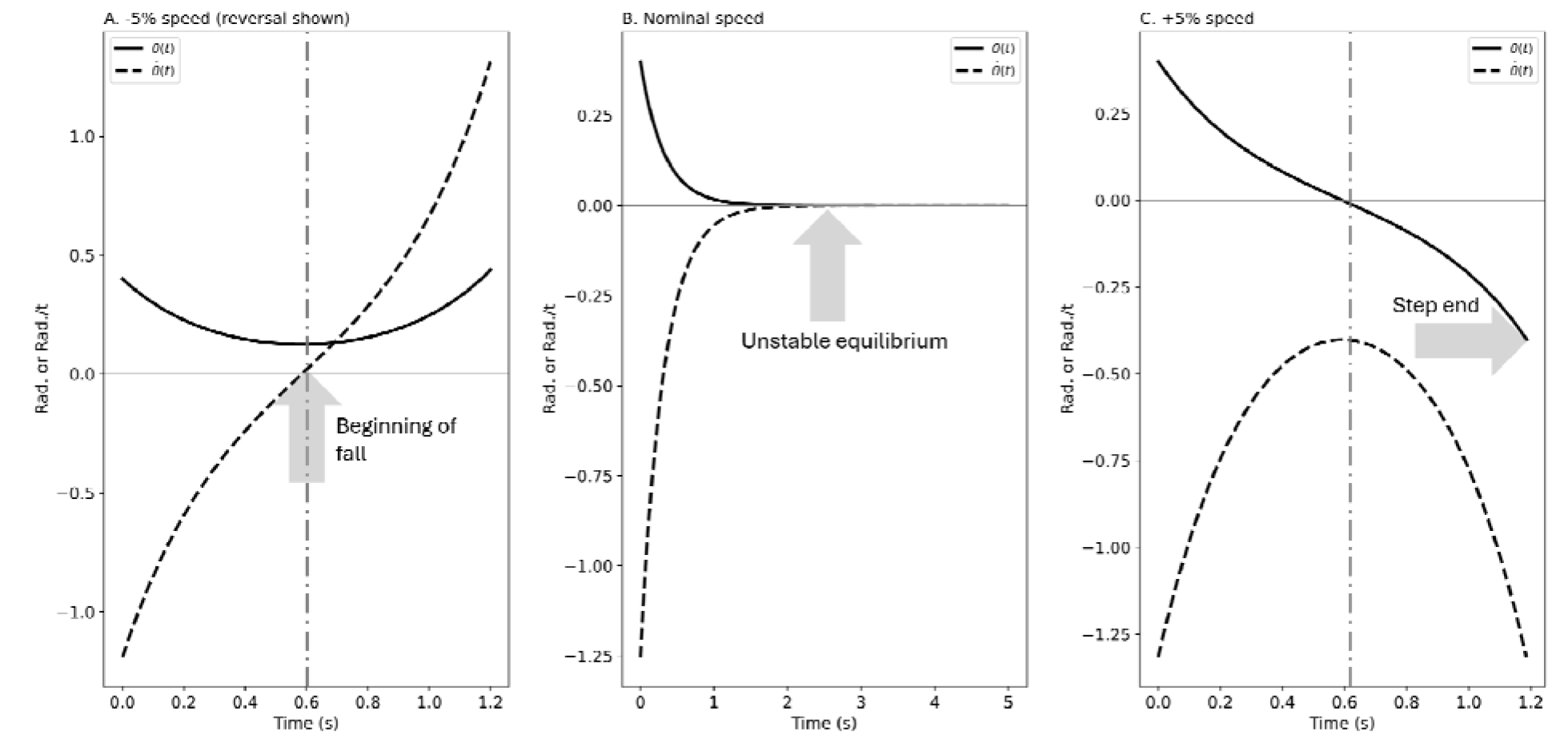
Based on the inverted pendulum equation of motion, sufficient initial kinetic energy is required to complete the gait. For a given step length, (A) when the initial COM speed is insufficient, the forward COM motion stops, and it begins to fall back. When COM speed becomes zero, the falling starts (B) When the initial speed is equal to the threshold, the COM reaches the mid-stance which is unstable equilibrium. At this point, a small perturbation may cause it to fall in either direction. (C) When the initial speed exceeds the threshold, the COM completes the gait.

**Figure 4.**
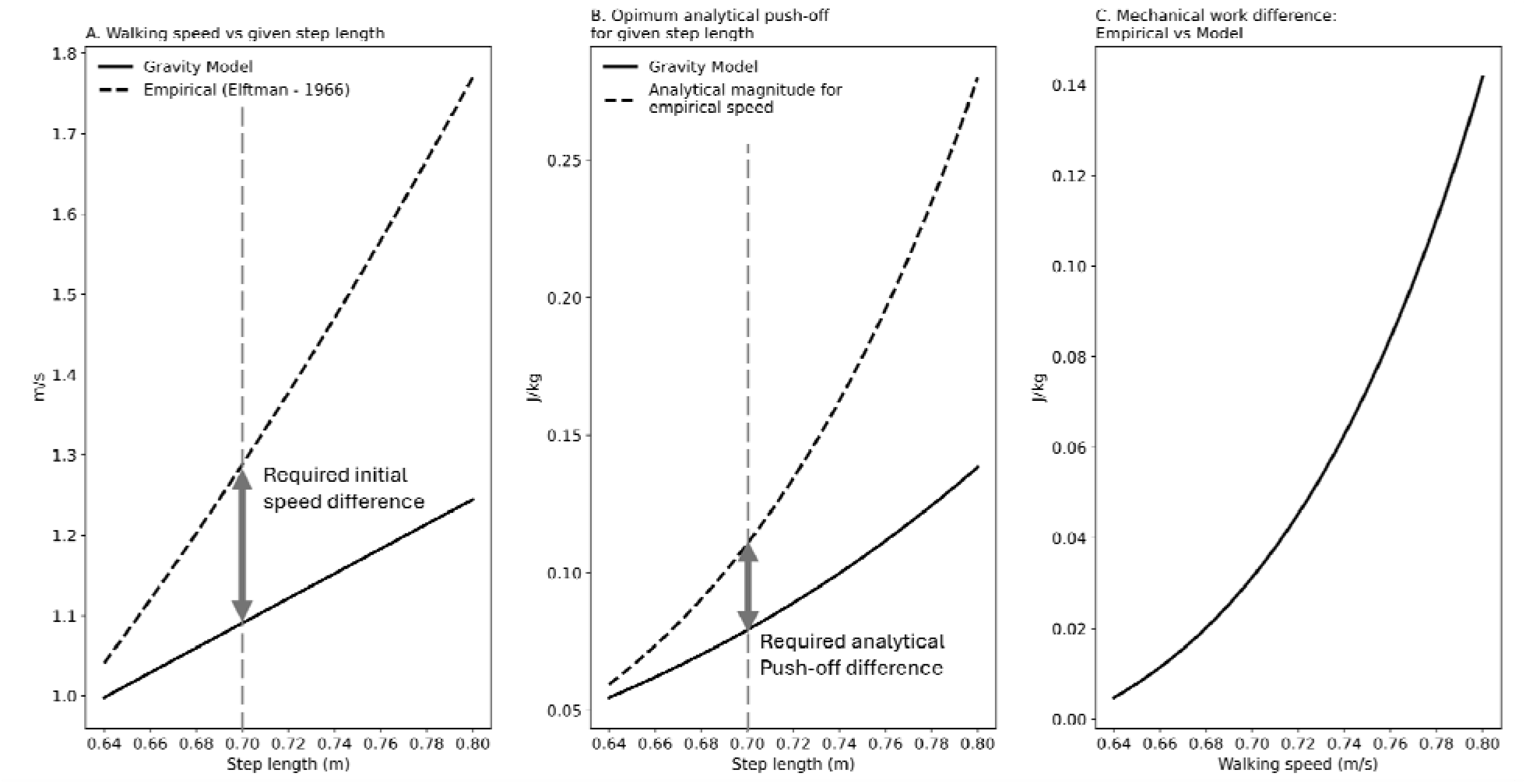
The magnitude of the single support gravity work rises with the increase of the step length prescribed, (A) to over come the gravity work associated with a given step length, the single support initial speed (COM speed after the step transition) must increase accordingly. (B) To restore the energy loss of the step-to-step transition, walker needs to preform positive mechanical work. The preferred method is to exert a pre-emptive push-off. With step length rise, step-transition work increases. Based on empirical step length and speed relationship, human walking requires larger walking speed. (C) The difference between step-transition work based on gravity method and empirical data must represent a form of energy dissipation in human walking.

We evaluate the work against gravity when the step length increases from 0.64 m to 0.8 m. The associated necessary walking speed ranges from 1.00 to 1.25 . Therefore, the associated analytical push-off increases from 0.05 to 0.14 . On the other hand, for prescribed step lengths, the empirical power law suggests a walking speed range of 1.04 to 1.77 . Therefore, the step-transition push-off varies from 0.06 to 0.28 .

That results in a step-transition work difference of 0.01 to 0.14 between the two methods. This means that the nominal analytical required push-off of real human walking is more than the gravity method suggests. Therefore, the human walking pendular phase must have some energy dissipation.

For S = 0.7 m, the gravity method suggests a minimum required walking speed of 1.1 . The pendular method (quadratic) indicates a maximum vertical reaction of 1.00 BW at midstance (Figure 5A). The linearly damped model, however, predicts an ‘M’ shape trajectory, but the average vertical force is 1.19 BW. Applying the muscle force adjustment trained by the experimental data (12), the,, and becomes -0.29, 0.09, and 0.75 respectively.

**Figure 5.**
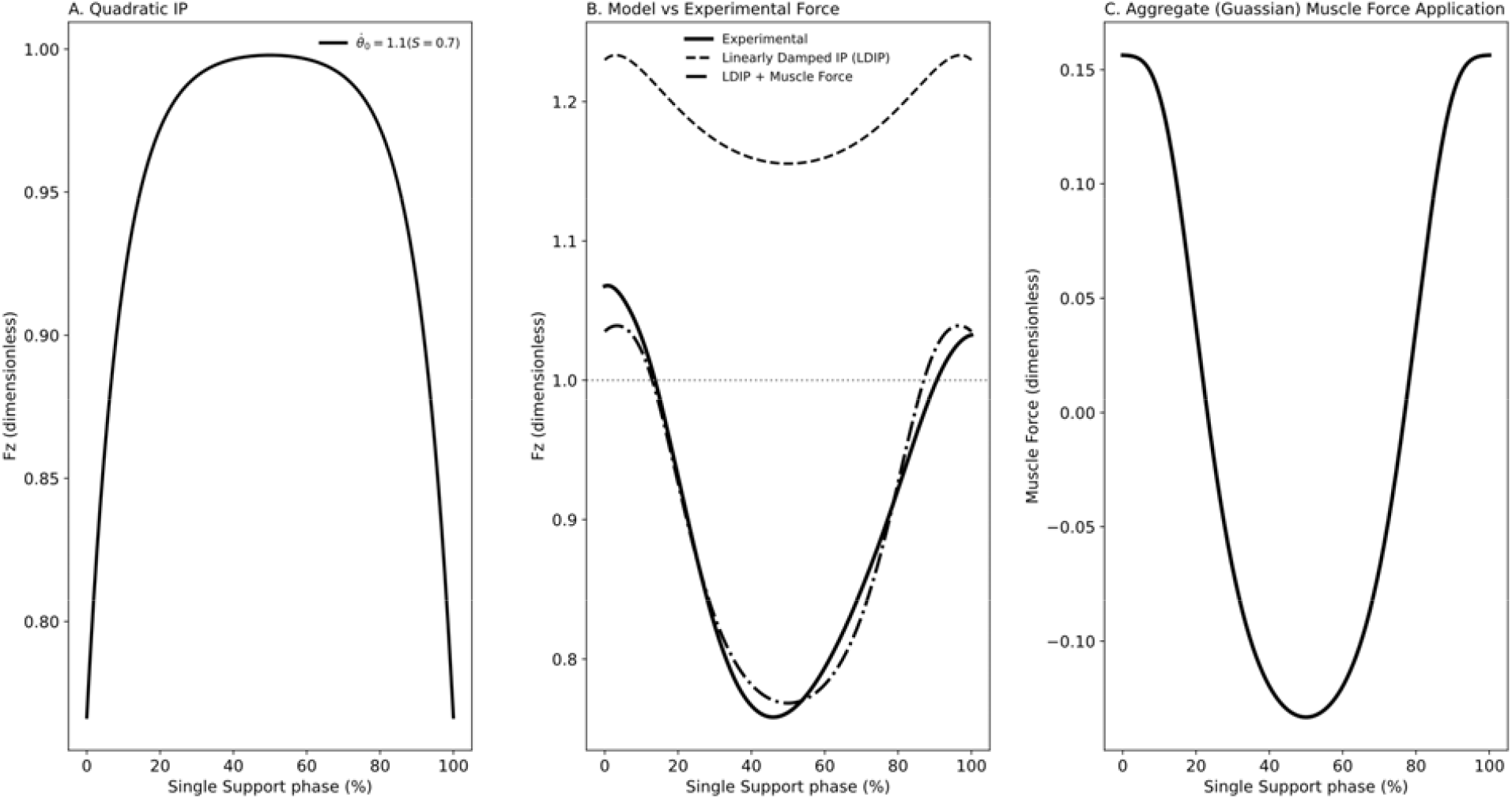
Analysis result for (A) pendular model with quadratic shape indicating maximum vertical force of 1.00BW, (B) linearly damped model without and with muscle intervention (gaussian) trained with experimental vertical GRF of the same step length (S = 0.7m), (C) muscle actuation (intervention) with net impulse of zero of the single support phase.

The resultant Gaussian trajectory demonstrates a vertical force that is like experimental vertical GRF during the single support phase (0.89BW) with a midstance trough of 0.77 BW (Figure 5B).

The muscle force suggests alternating forces during single support, positive at early and lat stance (+0.16BW) and negative at midstance (-0.13BW), indicating a net impulse of zero (0.00BW, Figure 5C).

When step length increases (S=0.6, 0.7, and 0.8m), the pendular model suggests that the maximum peak force is equal to 1.00BW, while the linearly damped model with muscle intervention (current model) suggests increasing peaks (collision and push-off) from 1.00BW to 1.04BW and 1.08BW respectively. On the other hand, midstance troughs also increase from 0.75BW to 0.77BW and 0.79BW respectively (Figure 6).

**Figure 6.**
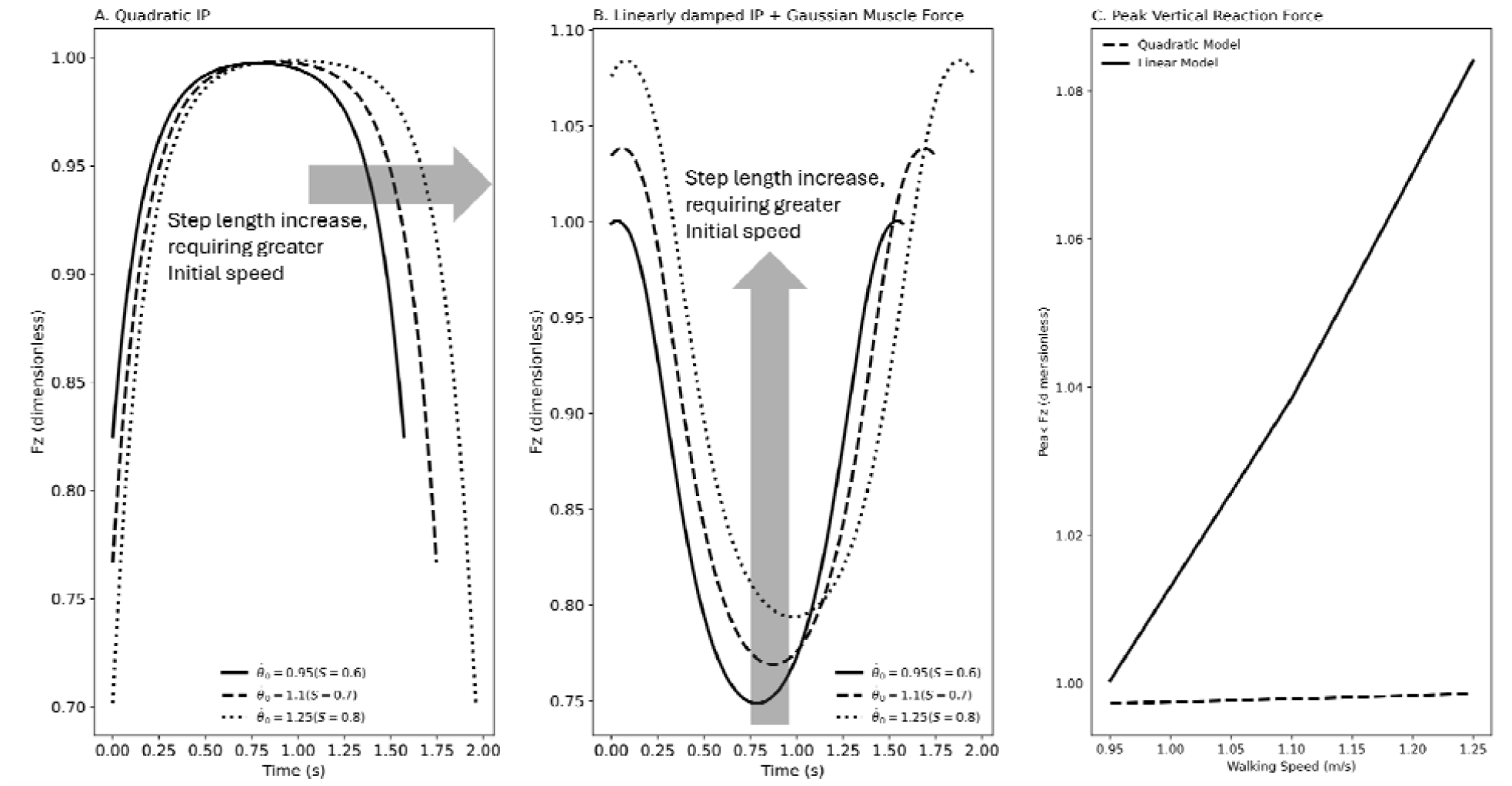
As the step length is increased, the gravity model indicates that greater initial speed is necessary to finish step. (A) Vertical reaction force of inverted pendulum represents a single vertex due to its quadratic nature. (B) The vertical reaction force of linearly damped model with muscle intervention (current model) depicts a typical ‘M’ profile similar to human walking. (C) The peak load of quadratic model changes marginally, while the current model suggests larger peak force with step length increase.

For a step length of S = 0.8 m, when walking speed increases from the minimum required by the gravity method (1.25) to 1.30, 1.35, and 1.40, the pendular model predict a decreasing trend in maximum vertical force, from 1.00 BW to 0.99, 0.97, and 0.96 BW, respectively. In contrast, the current model predicts an increase in peak vertical forces, from 1.08 BW at v = 1.25 m·s□^1^ to 1.08, 1.11, and 1.13 BW at the higher speeds, consistent with the shorter step durations at faster walking speeds (Figure 7).

**Figure 7.**
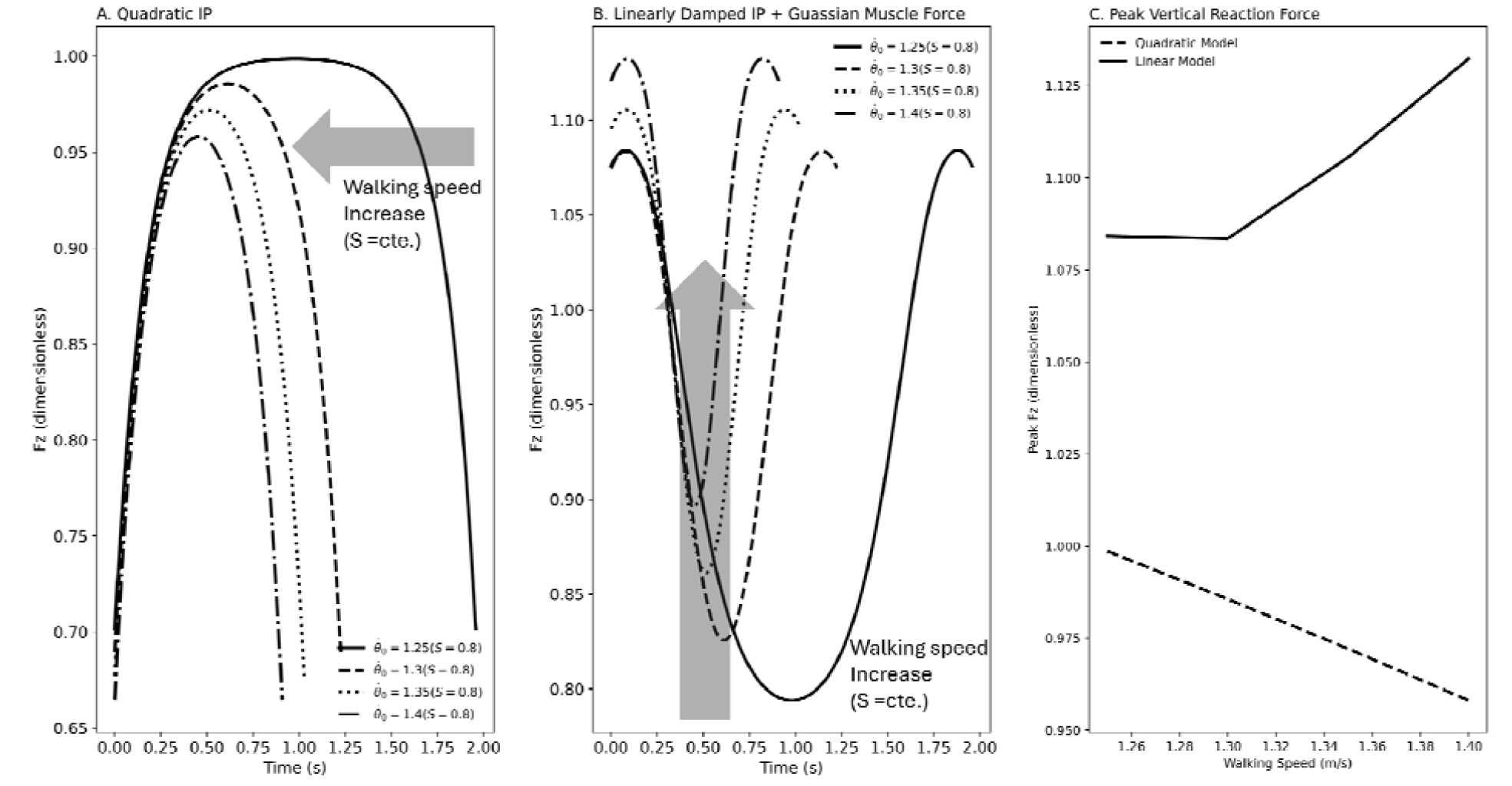
The vertical reaction force changes, when for a constant step length (S = 0.8m), the walking speed is increased from minimum required by gravity method. (A) Inverted pendulum model that has a quadratic trajectory, (B) current model indicates increasing collision and push-off peaks. (C) While the peak reaction force declines in inverted pendulum model due to growth of centripetal force, the current model suggests increase of force peaks with speed increase similar to human walking.

When (free motion at 0.4 rad and 1.25), the force is symmetrical and the trough (0.83BW) appears at the center of the trajectory. The pendular model maximum force is 0.98BW, while the current model’s collision and push-off peaks are 1.08BW with a step duration of 1.23 s. When (opposing), the profile and trough (0.80BW) are skewed to the right. While the pendular model predicts maximum force to be 0.99BW, the current model’s collision and push-off peaks become 1.11BW and 1.08BW respectively with a step duration of 1.86 s On the other hand, with supportive torque () the trajectory and trough (0.85BW) are skewed to the left. The pendular maximum vertical force is 0.97BW whereas, the current model predicts 1.08BW and 1.10BW for collision and push-off peaks with a step duration of 1.04 s (Figure *8*). Therefore, the single support active hip work impacts the amplitudes of both peaks and troughs.

**Figure 8.**
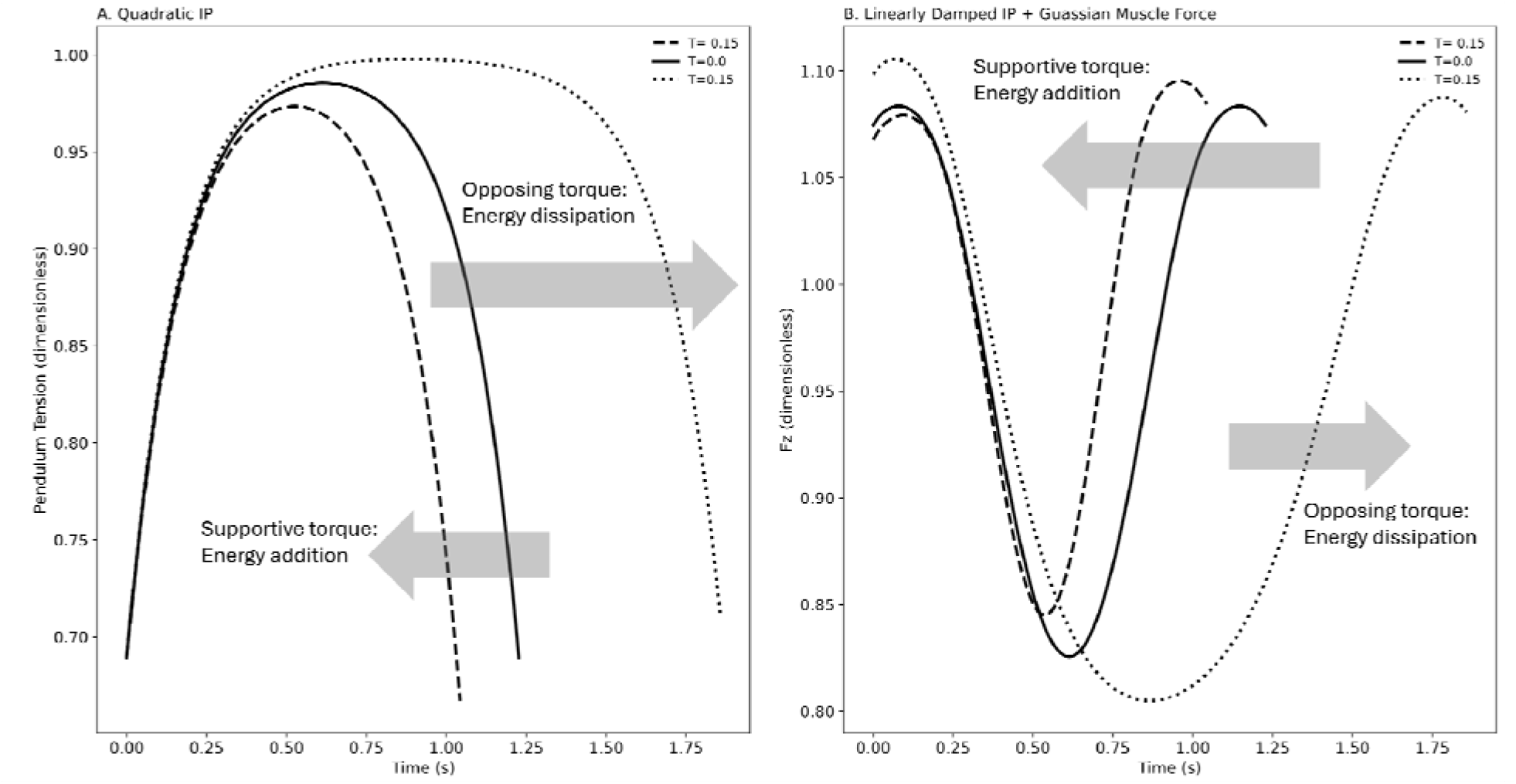
Application of a torque changes the energy of COM. If the torque is supportive, i.e. in direction of motion, the COM energy increases, and the step finishes faster with push-off peak larger than the collision peak. On the contrary, when the COM energy is wasted by an opposing torque, the push-off peak declines and it takes longer to finish the step. The COM motion with external torque application (A) inverted pendulum model, (B) linearly damped pendulum model with muscle intervention (current model).

Similar to human walking, we assume that the hip torque strategy changes after the step-to-step transition. We define the first vertical force peak as the onset of single support (15). After this point, the hip torque is held constant and negative (supporting). When hip torque is negative

(supporting) throughout the entire simulation, increasing 0 to 3.0 N.m.kg^−1^ the net hip work rises to 2.4 J.kg^−1^. When the torque is zero prior to the first vertical force peak, net hip work reaches 1.75 J.kg^−1^. When hip torque switches sign from positive (opposing) to negative (supporting) at the first peak, net hip work is 1.11 J.kg^−1^. Across all cases, the change in collision loss associated with the increased collision peak ranges from 0 to 0.13 J.kg^−1^.

Therefore, reversing the hip torque direction at the first peak yields the smallest net hip work while it still produces the observed peak differences. Step duration also decreases from 1.23 s to 0.49 s consistent with speed increase stemming from COM energy rise. When supportive hip torque is applied only from mid-stance onward, net hip work ranges from 0 to 1.12 J.kg^−1^, while step duration decreases from 1.23 s to 0.98 s (Figure 9).

**Figure 9.**
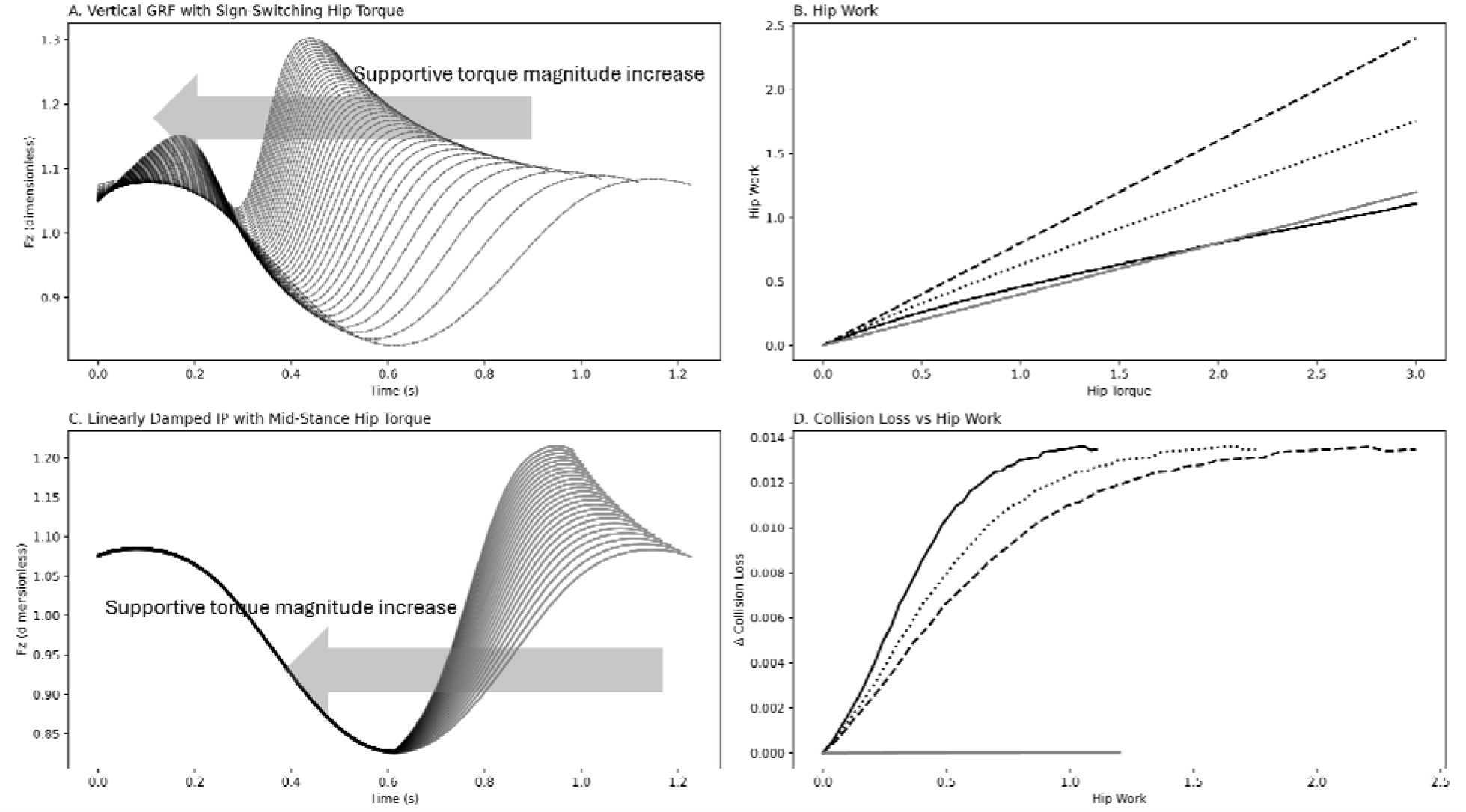
(A) As hip torque magnitude increases, the difference between the vertical force peaks increases, while step duration decreases. (B) Increasing hip torque also increases hip mechanical work, and (D) increases the change in collision energy loss. Switching hip torque from positive (opposing) to negative (supporting) after the first peak requires less hip mechanical work to achieve the same peak-amplitude differences than either applying zero torque before the first peak or applying negative (supportive) torque throughout the entire stance. Across torque strategies, the increase in collision loss is nearly the same. (C) When supportive hip torque is applied only from mid-stance onward, the collision peak does not change (i.e., collision loss remains constant). However, despite requiring nearly the same hip work as the torque-switching strategy, the resulting push-off peak difference is smaller. The solid line represents opposing hip torque up to the first peak, followed by supportive torque thereafter. The dotted and dashed lines represent zero torque and supportive torque applied before the collision peak, respectively. The gray line represents supportive hip torque applied from mid-stance onward.

## Discussion

This study demonstrates that preferred walking behavior is governed not only by energetic optimality (11) but also by mechanical feasibility constraints arising from step length, gravitational work, and step-to-step energy losses (6). While conservative inverted pendulum mechanics (2,4) define the minimum conditions under which a step can be completed, human walking systematically deviates from this ideal due to dissipation and active regulation of COM motion (10,17). By extending the pendular framework by linearizing its axial tension, structured force redistribution, and introduction of active work (hip), the present work provides a unified mechanical interpretation of walking speed selection, vertical ground reaction force (GRF) shape, and step transition work. Hence, this work attempts to cover a missing factor in human walking simulation. While the step-to-step transition active work is analytically quantified (5), the single support regulatory role in human gait done through active work is investigated in a limited number of studies (7–9,18).

The inverted pendulum model (2,3) remains valuable for identifying lower bounds on walking speed and momentum for a given step length. Our results reaffirm that, in the absence of sufficient initial kinetic energy, forward progression under those conditions become mechanically impossible. However, the discrepancy between gravity-based predictions and empirically observed walking behavior (10) indicates that human gait does not operate at this lower bound. Instead, humans walk with excess kinetic energy relative to conservative requirements, implying unavoidable dissipation also during single support (16,17).

Here, we indicate that single-support walking involves systematic active regulation of COM dynamics, even when net mechanical work over the step is minimal, i.e. walking at the mechanically preferred speed. Linearization of pendulum axial force captures first-order deviations from minimum momentum to meet the work against gravity demand, while the muscle modulation term redistributes vertical force without altering net impulse. Together, these features reconcile pendular mechanics with experimentally observed GRF profiles (19–21). Consequently, the damping-like term does not represent mechanical energy loss, but it redistributes forces within the stance phase and converts kinetic energy into temporary vertical support (early stance), controlled unloading (mid-stance), and finally, redirection into forward motion (late stance).

The parameters *C*_*Baseline*_, *A*_*m*_, and *σ*_*m*_ indicate that humans attempt to exploit pendular motion with minimum intervention while limiting walking load (22). *A*_*m*_ quantifies the magnitude of active force redistribution applied during stance, indicating how strongly muscle action reshapes vertical loading to manage gravitational and inertial demands. Having *A*_*m*_ = −0.29 indicates a substantial and likely purposeful force modulation to reduce vertical weight support (∼30% of BW). It indicates that muscles are allowing a controlled unloading by a slight downward COM acceleration (23). *σ*_*m*_ governs the timing of this redistribution, capturing whether muscle intervention is sharply localized or broadly distributed across single support. *σ*_*m*_=0.09 rad shows a highly localized muscle action that is concentrated about midstance. Therefore, muscle actuations must be precisely timed to provide a targeted intervention when passive dynamics are not completely favorable, but the intervention is minimal. The phase-localized muscle impedance modulation reproduces human-like vertical and horizontal ground reaction forces while maintaining near-zero net center-of-mass work across stance. As such, the current model also remains conservative through stance. Finally, *C*_Baseline_ characterizes the average vertical load borne by the stance limb during single support, reflecting the degree to which body weight support is distributed across phases of the step and between limbs. *C*_Baseline_∼0.79 indicates an attempt to carry less than full weight during single support, and to shift the peak forces to double support when greater vertical load is supported by both legs. Thus, *C*_Baseline_ must reflect inter-limb coordination and not muscle effort. Importantly, these parameters allow the shape of the vertical GRF to emerge from mechanistic principles rather than being prescribed a priori. This distinguishes the present framework from descriptive or purely empirical GRF models.

Human walking is work-stability optimized. From a control viewpoint, single support modulation is not a force tracing, stiffness maximization, nor a PID like correction, but it is a phase-based impedance shaping with minimal control to ensure optimal mechanical energy control. Hence, in single support, the muscle actions are to modulate the vertical GRF that comes about through changes in leg segments’ acceleration, geometry, or direction of forces (24–27). The model indicates a hierarchal low order control strategy with three layers. First is passive pendular motion (28,29) followed by a low-bandwidth modulation that is low dimensional and state dependant (*f* (*θ*)), i.e. two parameters of control only (*θ* and 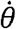). Thus, we may infer that this layer of control is a phase dependant (phase based) feedforward controller that reshapes the passive dynamics (impedance shaping) (30,31). The last layer is load-shaping that is slow and quasi-static for load sharing through inter-limb coordination (gain scheduling) (32). Therefore, it is fair to infer that single support control does not require explicit force feedback, nor deals with high gain error. Thus, it features predictive activation patterns and robustness to sensory noise (33–35).

The comparison of a different hip torque application strategy indicates the benefit of altering torque direction. That is, opposing in early stance and supporting for the rest of stance which is superior to hip torque of zero or supporting torque in early stance. While applying supporting hip torque from midstance adds more mechanical energy (+%9.1), it does not contribute to increase of collision loss (%1.2). Therefore, when the control system is able to regulate action timings, the hip energy infusion from mid-stance, similar to human walking (36), should be most favorable. Otherwise, alternative hip actuation from the beginning of single support presents a small penalty, while the duration of the step may be reduced substantially. It can be a practical strategy when the stance leg load support capacity or step-transition work capacity is limited (37,38). The other cases, i.e. supportive torque from the beginning or just after the collision peak, represent large mechanical energy losses. The energy loss stems from the forced constraints. Hip torque does not alter vertical or horizontal GRF in the current model because the rigid-leg depicts a single-DOF constraint that eliminates any actuator work that would change COM motion. The extra hip work is dissipated internally by the constraint, not by ground reaction forces. This may also be true for human walking as the COM motion trajectory is somewhat bounded by stance leg geometry. This energy loss indicates that if a walker requires the hip to cover a greater portion of energizing the gait to contribute to late stance push-off, i.e. beyond mechanical storage and release (6,7), hip supportive actuation must happen after midstance.

The present results clarify how step length and walking speed jointly influence force production. As step length increases, greater initial momentum is required to complete the pendular phase, which in turn elevates step transition work and peak forces (5,10). Similarly, increasing walking speed at fixed step length amplifies dissipation and shortens stance duration, necessitating higher force amplitudes to redirect the COM (8). Notably, the conservative pendular model predicts almost constant peak forces with increasing speed (2), whereas the extended model predicts material increases. This divergence highlights the critical role of active control in shaping external loading and underscores why passive models fail to capture observed force scaling in human walking (3).

The hip torque analysis further emphasizes that ***when*** work is applied it is as important as ***how much*** work is performed. Across conditions, altering hip torque timing had a larger effect on net hip work than altering peak force magnitudes. Minimizing hip work by switching torque direction near the onset of single support reinforces the notion of a just-in-time control strategy (33,39), in which energetic contributions are delivered at mechanically advantageous moments (40). Having supportive hip torque in early stance, nevertheless, enhances the collision peak and its associated loss undermining its contribution to the push-off rise. When hip work infusion is delayed until after midstance, while the magnitude of collision loss stays the same, the magnitude of exerted work becomes smaller yielding a smaller push-off. This result provides a mechanistic explanation for experimentally observed hip power patterns (36) proposing that the hip behaves more like an impedance modulator than a pure motor.

Hence, the analysis indicates that it is possible to have a pre-emptive push-off for the step transition with only hip actuation, but at elevated cost (41). In other words, when the ankle is eliminated, e.g. amputees, hip actuation, whether during single support or late stance increases the push-off peak (1,8,36) and elevates the COM energy (7). The model, however, does not impose any restriction on the magnitude of hip work. Nonetheless, there must be some physiological limits for hip work magnitude and its rate of application (42). Thus, we expect much lower push-off when only the hip contributes to COM energy. It is supported by experimental observation since it is reported that for different walking regimes, the contribution of ankle push-off is about 40%-60% of the step work while the hip contribution reaches only 20%-30% (43). The observed elevated cost of actuating the hip compared to the ankle may further support this viewpoint (41). As such, we expect a lower adopted walking speed for amputees and those with distal deficits (43). It is yet another support for the original idea of this study indicating that the capacity of positive work performance dominates other walking parameters such as adopted step length and associated walking speed. Therefore, one strategy to enhance walking mechanical energy is to increase elasticity to capture and release a portion of heel-strike collision loss (44) similar to the role of the Achilles tendon (45–47), while augmenting hip actuation with a simple exosuit setup inserting supportive positive work from midstance until toe-off.

In this study, several limitations should be acknowledged. First, the model assumes massless legs and rigid segments, neglecting segmental dynamics and joint-level energetics (5). We also restrict the joints’ actions to hip torque only. While this simplification enables analytical insight, it limits direct quantitative prediction of joint moments and muscle forces (43). Second, muscle action is represented phenomenologically through force redistribution rather than explicit muscle-tendon dynamics. Although this approach captures essential features of vertical GRF modulation, it does not distinguish between specific muscle groups or neural control strategies. Additionally, like the original pendular model (2), the linearized model is also conservative, thus, it does not capture nonlinear viscoelastic behavior, soft tissue wobbling, or foot–ground compliance (16,17). Future work should examine whether inclusion of dissipation terms improve model fidelity across a broader range of walking conditions as indicated by the difference between gravity work derived walking speed and empirical data. In this study we use step lengths and speeds that are at suggested mechanically preferred walking speed (S = 0.7 and associated speed *v*_*ave*_ = 1.2 m.s^−1^) (7), or rather close, in order to have small single support net active work. As such, for speeds or step lengths that are drastically different, *C*_Baseline_,*A*_*m*_, and *σ*_*m*_ the proposed magnitude may not hold. Presence of single support active work, whether dissipative or additive, may suggest an optimization to include parameters such as hip torque and timing to train the model. An increase in the mid-stance trough with speed is contrary to what is observed in human walking (48). We may associate the amplitude of the human walking trough to the active work performed during the single support deviating it from conservative. The influence of hip work on the reaction force trajectory is also noted in our analysis. Finally, the present study focuses on steady, level walking; extending the framework to uneven terrain, perturbations, and asymmetric gait will be necessary to fully test its generality.

Despite these limitations, the proposed framework offers a parsimonious yet mechanistically grounded extension of classical walking models. The modified pendular model complements the existing models that focus on the step-by-step transition (5) by depicting single support as an actively regulated phase. Although it presents a nominal GRF trajectory for given walking speeds, to represent a closer perspective of true human GRF, the force trajectory may be tuned by adjusting hip torque. By explicitly linking step length, walking speed, force redistribution, and work timing, it provides a foundation for future studies aimed at understanding and augmenting human locomotion. Together, these findings suggest that preferred walking speed emerges from an interaction between step length, mechanical work requirements, and the capacity to deliver that work efficiently. Rather than being selected solely to minimize metabolic cost, walking speed must satisfy minimum momentum and work constraints while limiting the need for energetically unfavorable compensatory strategies. This framework has implications for understanding gait adaptations in populations with reduced work capacity, such as older adults or individuals with neuromuscular impairments, as well as for the design of simple but effective assistive devices (49,50) that aim to restore or augment step-to-step work.

## Acknowledgment

This work was supported by a Natural Sciences and Engineering Research Council of Canada (NSERC) Discovery grant (04823-2017) received by J.E.A.B.

## Notes

**Conflict of Interest:** None

### Competing Interest Statement

The authors have declared no competing interest.

